# Primary visual cortex injury produces loss of inhibitory neurons and long-term visual circuit dysfunction

**DOI:** 10.1101/2020.08.19.258335

**Authors:** Jan C. Frankowski, Andrzej T. Foik, Jiana R. Machhor, David C. Lyon, Robert F. Hunt

## Abstract

Primary sensory areas of the mammalian neocortex have a remarkable degree of plasticity, allowing neural circuits to adapt to dynamic environments. However, little is known about the effect of traumatic brain injury on visual system function. Here we applied a mild focal contusion injury to primary visual cortex (V1) in adult mice. We found that, although V1 was largely intact in brain-injured mice, there was a reduction in the number of inhibitory interneurons that extended into deep cortical layers. In general, we found a preferential reduction of interneurons located in superficial layers, near the impact site, while interneurons positioned in deeper layers were better preserved. Three months after injury, V1 neurons showed dramatically reduced responses to visual stimuli and weaker orientation selectivity and tuning, consistent with the loss of cortical inhibition. Our results demonstrate that V1 neurons no longer robustly and stably encode visual input following a mild traumatic injury.

**Highlights:**

- Inhibitory neurons are lost throughout brain injured visual cortex
- Visually-evoked potentials are severely degraded after injury
- Injured V1 neurons show weaker selectivity and tuning consistent with reduced interneurons

## Introduction

In primary visual cortex (V1), GABAergic inhibition is essential for several basic functions, such as tuning a neuron’s preference for stimulus contrast, size and orientation (Sillito et al., 1985; Cardin et al., 2007; Lee et al., 2012) as well as higher-order processing, such as contrast perception (Cone et al., 2019). During development, cortical inhibition modulates critical periods, a transient time of enhanced sensitivity to sensory experience. This has been most extensively studied in juvenile V1, in which obstructing vision through one eye results in cortical blindness to this eye, even after normal vision is restored (Wiesel, 1982). Cortical inhibition is required for opening the developmental critical period in visual cortex (Hensch et al., 1998) and inactivating interneurons can prolong the critical period (Espinosa and Stryker, 2012) or impair cortical plasticity (Fagiolini et al., 2004). Even in adulthood, after binocular vision is well established, manipulating inhibition through pharmacology (Sillito et al., 1985; Fagiolini and Hensch, 2000) or interneuron transplantation (Davis et al., 2015; Larminer et al., 2016) can have dramatic effects on cortical plasticity in response to monocular visual deprivation. Lesions also trigger cortical plasticity and functional disturbances in visual cortex (Girard et al., 1991; Eysel et al., 1999; Imbrosci et al., 2010). However, no study has investigated the effect of traumatic brain injury (TBI) on visual circuit function *in vivo*.

Posterior impact injuries to occipital cortex are extremely common in human, and TBI can lead to long-lasting visual impairments, such as visual acuity and field loss, binocular dysfunction and spatial perceptual deficits (Sano et al., 1967; Stelmack et al., 2009; Armstrong, 2018). Following TBI in human, histological studies have documented a reduction in the number of GABA-producing interneurons in hippocampus (Swartz et al., 2006) and neocortex (Buriticá et al., 2009). In rodent models, TBI produces region-specific reductions in interneurons in various brain regions (Lowenstein et al., 1992; Toth et al., 1997; Santhakumar et al., 2000; Cantu et al., 2014; Huusko et al., 2015; Frankowski et al., 2018; Nichols et al., 2018; Vascak et al., 2018), and a loss of inhibition that does not recover with time (Toth et al., 1997; Li and Prince, 2002; Witgen et al., 2005; Hunt et al., 2011; Pavlov et al., 2011; Gupta et al., 2012; Butler et al., 2016; Almeida-Suhett et al., 2014, 2015; Vascak et al., 2018; Koenig et al., 2019). However, nearly all of the information about cellular responses to TBI in neocortex comes from studies evaluating somatosensory, motor or frontal cortex. A deeper understanding of functional disturbances in the traumatically injured visual cortex is important, because it has potential to provide a rational basis for therapy. To investigate central visual system trauma, we applied a focal controlled cortical impact (CCI) injury to produce a mild posterior impact TBI in mice. Then, we measured interneuron density and visually-evoked responses in V1 at chronic time points post injury.

## Results

### Occipital CCI produces a mild contusion in V1

We first evaluated the effect of a single, mild contusion injury in V1. To do this, we delivered mild CCI injury centered over the rostral end of V1 in young-adult GAD67-GFP reporter mice at P60 (Fig 1A, B). We selected CCI as a model, because the injury is highly reproducible from animal to animal, reliably recapitulates structural and functional deficits of TBI and focal contusion injuries are among the most common posterior impact injuries observed in human (Sano et al., 1967; Stelmack et al., 2009; Armstrong, 2018). In all CCI-injured animals (n = 6 mice), the lesion consisted of mild tissue compression that was restricted to superficial layers of the cortex at the injury epicenter (Fig 1C). At 14d post-CCI, a time point when lesion volume is largely stable (Hall et al., 2005; Pleasant et al., 2011), there was no significant difference in cortical volume between uninjured control and CCI-injured littermates (CCI: 95 ± 3%, compared to 99 ± 1% in control; P= 0.25; Fig 1D). However, when we evaluated the thickness of cortical tissue remaining at the injury epicenter in the most severely contused portion of the section, we found a 20% decrease in cortical thickness (CCI: 699 ± 46 μm, Uninjured: 880 ± 22 μm, P= 0.011; Fig 1E). This difference was only observed at the injury epicenter (0μm); no difference in cortical thickness was observed in tissue sections 300μm and 600μm caudal to the epicenter (Fig 1F). Thus, CCI produced a mild injury with minimal structural damage to V1.

**Figure 1.**
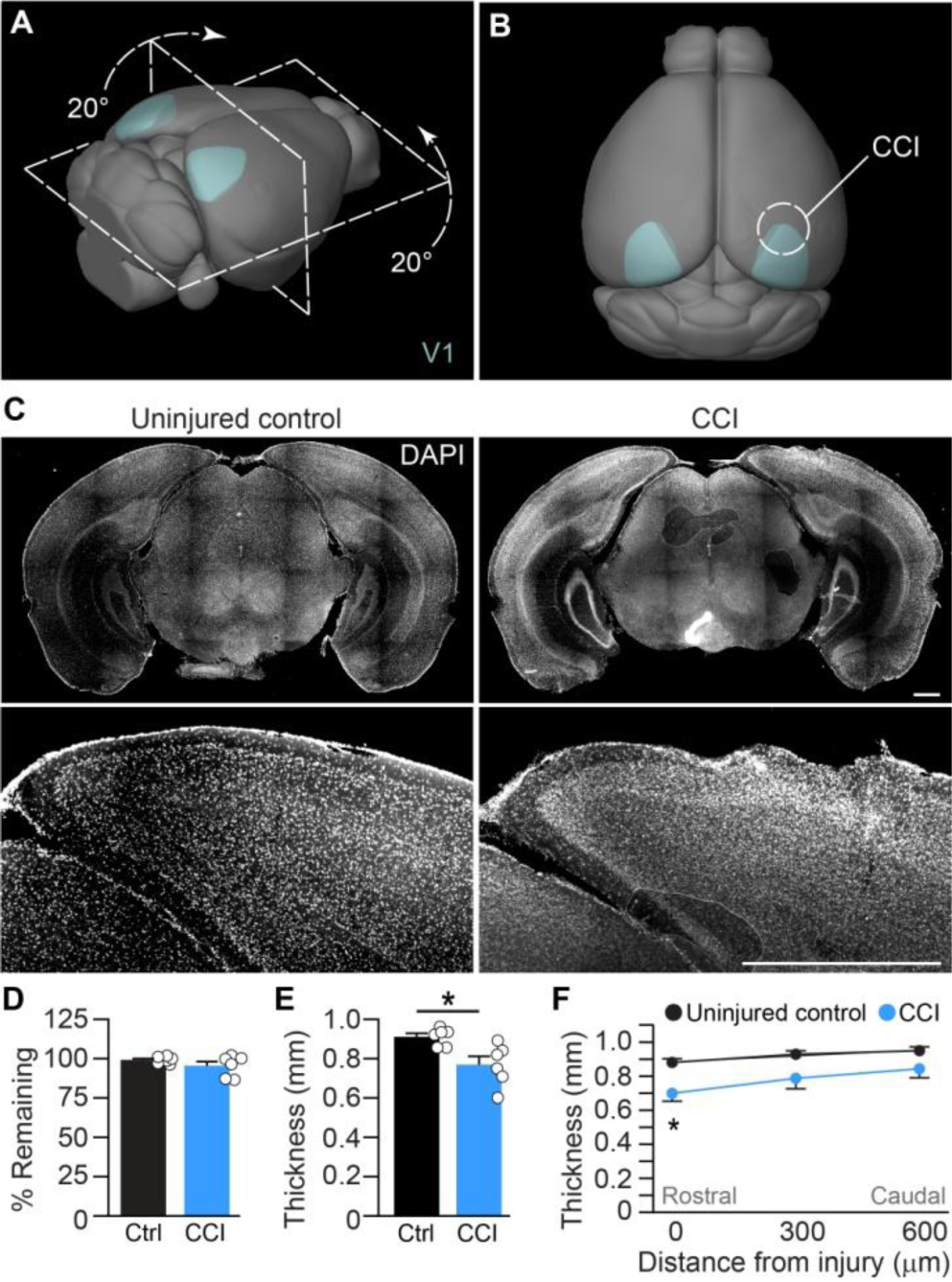
Visual cortex TBI produces a mild cortical lesion. **A.** Schematic of Allen Mouse Common Coordinate Framework showing the head rotation used to produce CCI injury over V1 (cyan). **B.** CCI injury was centered over rostral primary visual cortex (white circle). **C.** Coronal sections of DAPI labeling in a control animal and 14 days after CCI injury. **D.** Quantification of cortical tissue volume in control and CCI-injured litermates. **E.** Average thickness of cortex. *P = 0.012, two-tailed t-test, n = 6 mice / group. **F.** Average thickness of cortex with distance from the injury. *P = 0.011, Two-way repeated-measures ANOVA with Sidak post-hoc test, n = 6 mice per group. Scale bars, 1mm; error bars, SEM.

### Inhibitory neuron loss after V1 injury

Inhibition is critical for a wide range of V1 functions (Sillito et al., 1985; Cardin et al., 2007; Lee et al., 2012; Cone et al., 2019), and interneuron loss is a common feature of TBI in rodents and humans (Hunt et al., 2013). Therefore, we next quantified interneuron density in V1 using the same GAD67-GFP reporter mice. At 14d after CCI, we found ~20% decrease in the density of GAD67-GFP+ cells in V1 ipsilateral to the injury (Fig 2A; **Table S1**). No change in interneuron density was observed in the contralateral hemisphere. We also found that GFP+ cell loss was most severe at the injury epicenter and decreased with distance from the injury site (Fig 2B), similar to severe CCI injuries that damage hippocampus (Frankowski et al., 2019). To determine if post-traumatic interneuron loss was layer-specific, we quantified interneuron density in cortical layers 1, 2/3, 4, and 5/6 of CCI-injured and uninjured control littermates (Fig 2C-H; **Table S1**). We found that cell loss extended throughout the cortical column ipsilateral to the injury, with significant reductions in GFP+ cells in cortical layers 1, 2/3 and 5/6, but no interneuron loss was observed in layer 4. No change in GFP+ cell density was observed in any layer of the contralateral hemisphere.

**Figure 2.**
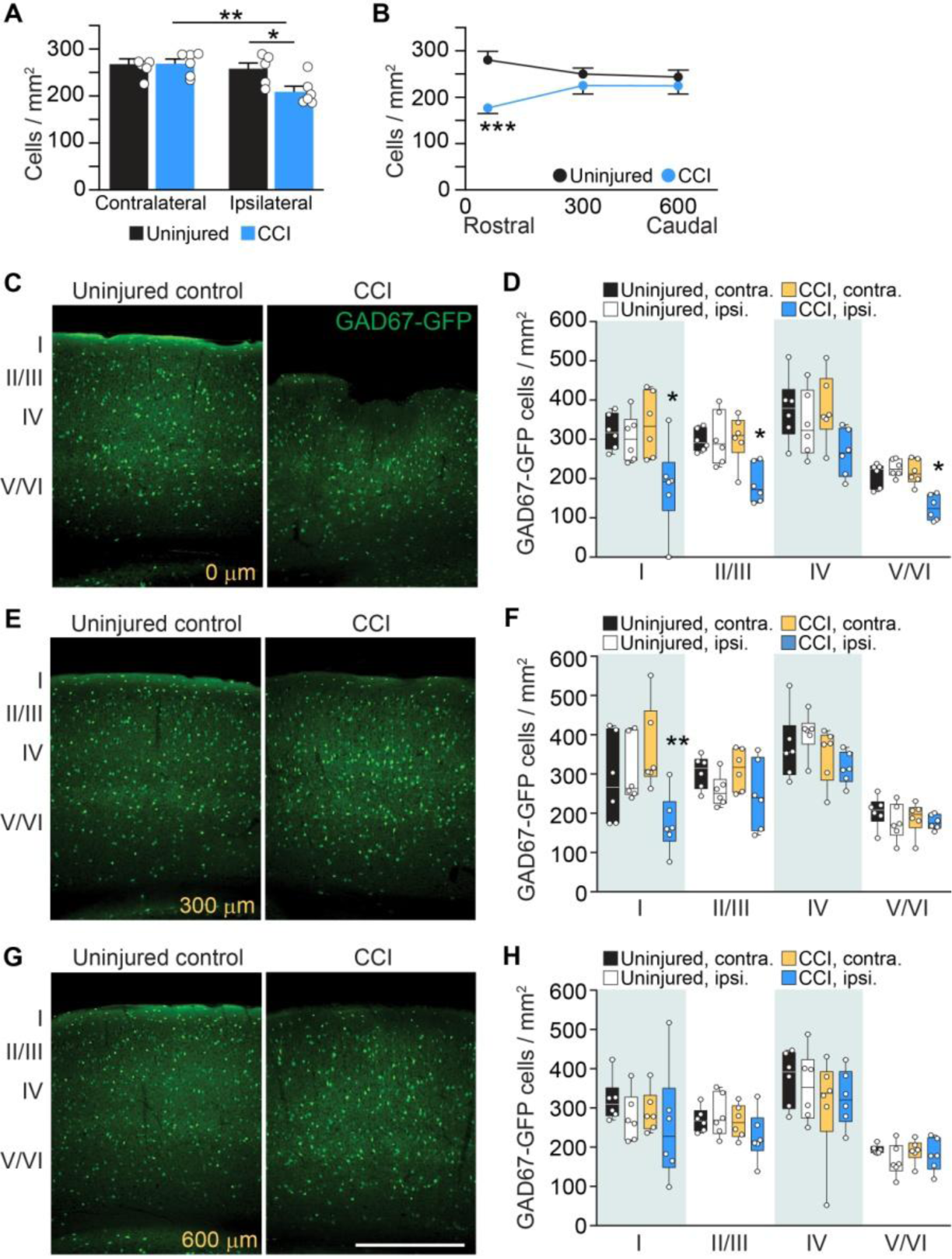
Interneuron loss in V1 after TBI. **A, B.** Quantification of GAD67-GFP+ cell density in control and brain-injured mice 14d after CCI. *P=0.0289; **P=0.0067; ***P = 0.0001, Two-way repeated-measures ANOVA with Sidak post-hoc test; n= 6 mice per group. **C, E, G.** Coronal images of V1 with GFP+ interneurons labeled in uninjured-control and CCI injury groups. **D, F, H.** Quantification of GFP+ cell density in layers 1, 2/3, and 5/6. *P= 0.0124, Layer I, injury site (D), *P= 0.0178, Layer III, injury site (D), *P= 0.0484, Layer V/VI, injury site (D), **P 0.0018 Layer I, 300um (F), Two-way repeated-measures ANOVA with Sidak post-hoc test; n= 6 mice per group. Scale bar, 500μm; error bars, SEM.

### Visually evoked responses in V1 are disrupted by brain injury

Although central visual system TBI is extremely common, and as many as 75% of military Service members suffer from visual dysfunction or functional blindness (Stelmack et al., 2009), there is essentially nothing known about visual circuit function after TBI. To evaluate the functional state of visual cortex following CCI, we measured visually evoked potentials (VEP) and single unit responses to a range of stimuli across a wide extent of injured V1 at 3 months after injury (Fig 3A). First, VEPs were recorded in response to brief flashes of light. Representative examples of flash-evoked responses are shown in individual animals (Fig 3B), along with population averages (Fig 3C). Compared to uninjured controls, evoked amplitudes were reduced significantly in brain injured mice, by more than 80% (Control: 3.2 ± 0.3 μV, compared to 1.4 ± 0.2 μV after CCI; P = 0.0003; Fig 3D), and response latencies were more than 60% longer (Control: 62 ± 4 ms, compared to 102 ± 10 ms after CCI; P = 0.001; Fig 3E). Of note, we also found that VEP amplitude reduced similarly across the site of injury, and wave profiles lacked a negative wave component normally present in deeper cortical layers (Fig S1).

**Figure 3.**
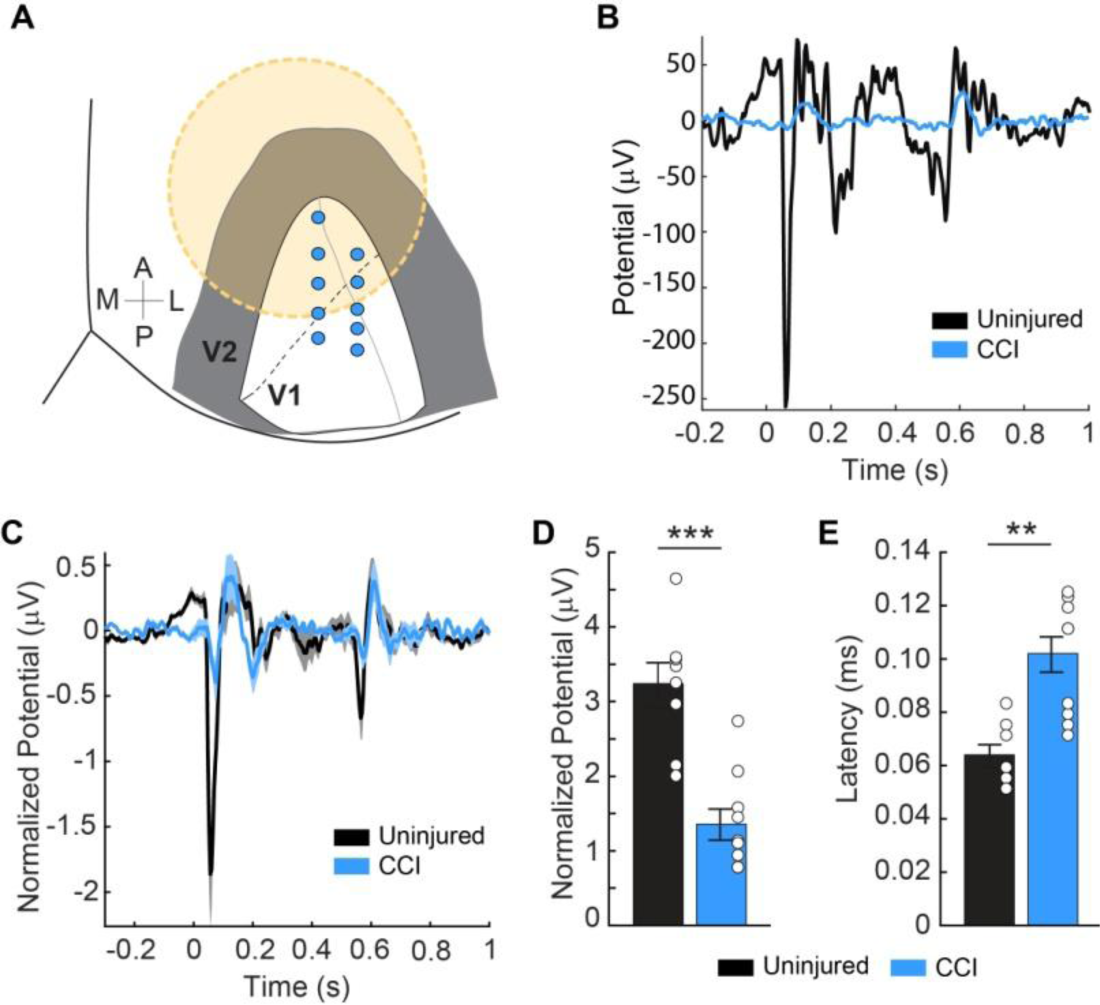
Visually evoked potentials are altered by TBI. **A.** Schematic showing the location of electrode placement in V1 (blue circles) relative to CCI injury (orange circle, 3 mm in diameter). **B.** Evoked potentials from a single uninjured control animal (black trace) and an animal 3 months after CCI (blue trace). **C.** Average evoked potentials from 10 recording locations per brain injured animal (light blue) and 9 sites per uninjured control mice. **D.** Quantification of average evoked amplitude. ***P=0.0003, two-tailed Mann-Whitney U, n= 9 locations (control), n= 10 locations (brain injury). **E.** Quantification of average response latency. **P=0.001, two-tailed Mann-Whitney U. Individual data points represent the value for each of the recording locations. Error bars, SEM.

To evaluate the functional profile of injured V1 in more detail, we next measured single neuron responses to a range of fundamental stimuli, including orientation, size, spatial frequency and temporal frequency. Representative examples of response profiles in Figure 4 show weaker tuning and selectivity to all four types of stimulus parameters in brain injured mice (Fig 4B,E,H,K) compared to uninjured controls (Fig 4A,D,G,J). For the cell population, these differences were significant for orientation (Fig 4C), size (Fig 4F) and spatial frequency (Fig 4I), but not temporal frequency (Fig 4L). The difference was quite striking for orientation and size tuning, both of which are strongly mediated through local cortical inhibition (Liu et al., 2011; Liu et al., 2015; Liu et al., 2017). For orientation, the tuning width (HWHH) of the preferred orientation is more than twice as sharp in the control example (24.3º vs 48.9º; Fig 4A,B), and this difference is also seen for the population (Control: 27.7º ± 2.4, compared to 47.9º ± 4.4 in CCI injured animals; P = 0.0009; Fig 4C). Broader tuning after TBI is consistent with orientation tuning mediated more through feedforward mechanisms and impairments in cortical inhibition (Liu et al., 2015; Liu et al., 2017). Similarly, the larger size preference in TBI compared to control neuron examples (73º vs. 35º; Fig 4D,E) and populations (52 ± 5 º, compared to 32 ± 3 º; P = 0.005; Fig 4F) also indicates a loss of cortical inhibition (Liu et al., 2011).

**Figure 4.**
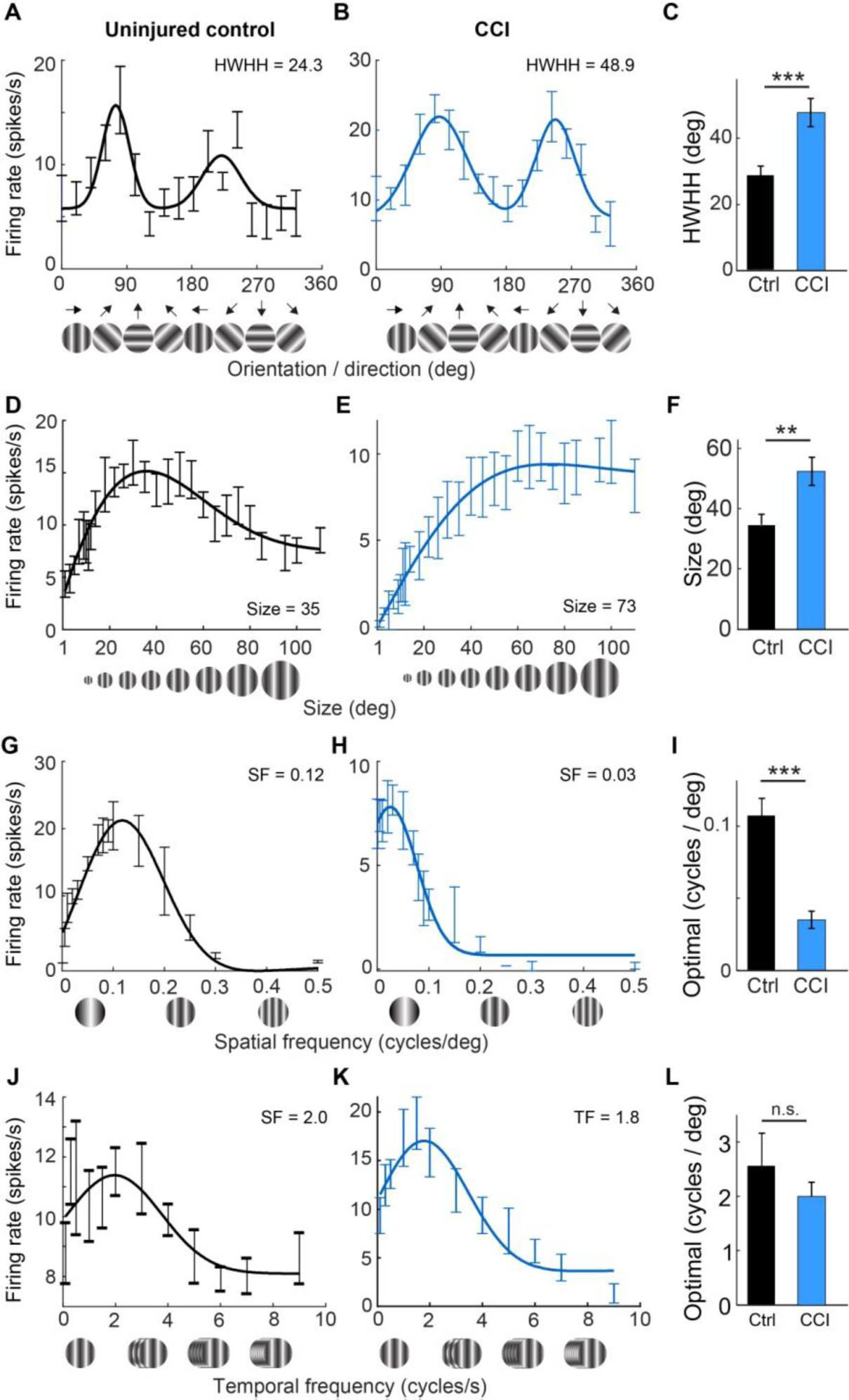
CCI disrupts V1 neuron tuning curves in response to drifting gratings. **A-C.** Orientation tuning curves for single neurons in an uninjured control (A) and CCI-injured mouse (B). HWHH tuning values are given in each panel and the population averages are quantified in (C). *** P = 0.0009, two-tailed Mann-Whitney U, n= 63 cells from 3 controls, n 39 cells from 3 brain injured mice. **D-F.** Single neuron examples and population average quantification of size. **P = 0.005, two-tailed Mann-Whitney U. **G-I.** Single neuron examples and quantification of spatial frequency (SF). ***P = 0.000004, two-tailed Mann-Whitney U. **J-L.** Single neuron examples and quantification of temporal frequency (TF). n.s., P = 0.15, two-tailed Mann-Whitney U. Optimal values for each parameter are given in each panel. Error bars, SEM.

## Discussion

Patients with TBI can show long-lasting deficits in visual system function, such as visual acuity and field loss, binocular dysfunction and spatial perceptual deficits (Armstrong, 2018). Here, we delivered a mild focal contusion injury directly to V1 to model occipital contusion injuries, which occur almost exclusively after a direct blow to the back of the head (Sano et al., 1967; Ommaya et al., 1971). We found that mild V1 injury produced inhibitory interneuron loss and long-lasting changes in visual circuit function. Although V1 was relatively well-preserved, compared to traditional approaches that produce substantial tissue loss (Frankowski et al., 2019), we found interneuron loss at the injury site that extended into deep cortical layers. *In vivo* recordings revealed a massive loss of visually evoked potentials and dramatically altered tuning to visual stimuli, including orientation and size, which have been shown to be modulated by cortical interneurons (Lee et al., 2012). These findings are consistent with human studies showing visual field defects can occur in individuals with no measurable lesion (Silverman et al., 1993).

Prior studies using blast injury have documented deficits in visual acuity and contrast sensitivity (Reiner et al., 2014). However, the diffuse nature of blast injury makes it difficult to determine whether visual deficits were due to central visual system injury or traumatic injury in the periphery. While chemical or laser lesions produce a more focal injury in V1 (Eysel and Schweigart, 1999; Imbrosci et al., 2010), these injury paradigms do not fully recapitulate mechanisms of TBI, which involves a mechanical force that disrupts brain function (Mendelow and Crawford, 1997). As such, our results represent the first demonstration of visual circuit changes following central visual system TBI.

The laminar organization of visual cortex and functional specialization of each cortical layer in sensory processing is generally conserved across species (Callaway, 1998). Layer 4 receives the majority of input arising from the lateral geniculate nucleus (LGN) and projects to neurons in layer 2/3, which in turn projects to deeper layers and other cortical areas (Miller et al., 2001). Layer 4 also plays an important role in generating response properties, such as orientation and direction selectivity (Zhuang et al., 2013). We found that layer 4 interneurons were not decreased in number after CCI injury. This may indicate that the first level of cortical processing in V1 could be preserved in these animals. In addition to visual input, V1 is a major relay for sensory information between other brain regions (Wandell and Smirnakis, 2009). For example, layer 1 contains a unique population of interneurons that receive sensory and neuromodulatory inputs, and they are responsive to sensory input beyond vision. These inhibitory neurons receive cholinergic input from the basal forebrain, which modulates the excitability of thalamocortical synapses in layer 4 (Alitto and Dan, 2012). Layer 1 also receives input from other sensory areas, such as primary auditory cortex (Ibrahim et al., 2016), and these neurons are strongly activated by non-visual input, such as locomotion, noise, and whisker stimulation (Mesik et al., 2019). Thus, interneurons located in layer 1 are involved in multi-modal sensory processing across different behavioral states. Based on our results demonstrating that layer 1 inhibitory neurons were most affected by visual cortex trauma, as compared to other cell layers, it is possible that the coordination of sensory information between different cortical areas may be profoundly disrupted in brain injured animals.

Individuals with TBI can develop visual impairments independent from other injury-induced motor or cognitive deficits (Zihl and Kerkhoff., 1990; Padula et al., 1994; Du et al., 2005; McKenna et al., 2006). Increases in light intensity evoke inhibitory synaptic activity to prevent changes in luminance intensity from disrupting cortical circuit function (Tucker and Fitzpatrick, 2006) and inability to modulate cortical gain has been proposed as a potential mechanism of injury-related photosensitivity (Zihl and Kerkhoff, 1990; Du et al., 2005). Here we show that basic visual processes in V1 are altered to reflect a distinct lack of cortically mediated inhibition. We found significantly broader orientation tuning widths consistent with reduced local inhibitory neuron activity (Lee et al., 2012). Instead, in brain injured animals, V1 orientation tuning resembles the broader widths mediated through feedforward mechanisms from the thalamus (Liu et al., 2015; Liu et al., 2017), which are likely more intact. Similarly, increased spatial summation indicated by larger stimulus size preference in TBI is consistent with the loss of local inhibitory neurons mediating surround suppression (Adesnick et al., 2012) and likely reflects preservation of feedforward mediated mechanisms (Liu et al., 2011).

## Author Contributions

J.C.F contributed to the execution and analysis of experiments and wrote the manuscript. A.T.F performed neurophysiology experiments and analyzed data. J.R.M. analyzed cell quantifications. D.C.L designed neurophysiology experiments, analyzed data, contributed funding and edited the manuscript. R.F.H contributed to the concept, design, analysis of experiments, funding and edited the manuscript.

## Acknowledgements

This work was supported by funding from the National Institutes of Health grants R01–NS096012, F31– NS106806, and R01–EY024890. We thank members of the Hunt lab for helpful discussions and comments on earlier versions of this manuscript.

## Methods

### Contact for Reagent and Resource Sharing

Further information and requests for resources and reagents should be directed to and will be fulfilled by the corresponding author, Robert F Hunt.

### Experimental Model and Subject Details

#### Animals

Mice were maintained in standard housing conditions on a 12h light/dark cycle with food and water provided *ad libitum*. All protocols and procedures followed the guidelines of the University Laboratory Animal Resources at the University of California, Irvine and adhered to National Institutes of Health Guidelines for the Care and Use of Laboratory Animals. All core and CCI-specific Common Data Elements (CDEs) defined by the Preclinical TBI Working Groups and the NIH/NINDS/DOD CDE Team (https://fitbir.nih.gov/content/preclinical-common-data-elements) can be found in **Tables S2** and **S3**. For electrophysiology experiments, we used CD-1 mice (Charles River, cat no. 022), and for anatomy experiments, we used a hemizygous glutamic acid decarboxylase - enhanced green fluorescence protein (GAD67-GFP) knock-in line (Tamamaki et al., 2003) maintained on a CD-1 background for > 10 generations.

### Method Details

#### Experimental design

Male and female mice were randomly allocated to experimental groups prior to TBI. Brain injury was performed at P60 and experiments were performed 14d (immunostaining) or 3 months (electrophysiology) after TBI. No data or animals were excluded from analysis.

#### Controlled cortical impact (CCI)

Unilateral controlled cortical impact was performed as previously described (Frankowski et al., 2019; Zhu et al., 2019), with modifications to the location and depth of injury. Mice were anesthetized with 2% isoflurane until unresponsive to toe-pinch, then placed into a stereotactic frame and maintained on 1% isoflurane. The fur overlying the skull was trimmed and the scalp was scrubbed with betadine before exposing the skull with a midline incision. The skull was rotated 20 degrees counterclockwise along the rostral-caudal axis and the rostral end of the skull was lowered 20 degrees relative to skull-flat. This orientation centered the impactor tip at the rostral end of V1 (Fig 1B). A ~4-5 mm craniotomy was centered 3 mm lateral to midline and 3 mm rostral to the lambdoid suture in the right hemisphere. The skull cap was removed leaving the dura intact. A computer-controlled pneumatically driven impactor (TBI-0310, Precision Systems and Instrumentation) with a 3 mm beveled stainless-steel tip was used to deliver a 0.2 mm depth contusive injury perpendicular to the dura at 3.5m/s velocity and 500ms of impactor dwell time. The skull cap was not replaced, and the incision was closed with silk sutures. Animals undergoing surgical procedures received buprenorphine hydrochloride (Buprenex, 0.05mg/kg, delivered i.p.) pre-operatively and once daily for 3d. A post-operative health assessment was performed for 5d following surgical procedures.

#### Immunostaining

Fourteen days after injury, mice were transcardially perfused with 4% paraformaldehyde (PFA) and free-floating vibratome sections (50 μm) were processed using standard immunostaining procedures (Zhu et al., 2019). Sections were stained with primary (GFP, 1:1000) and secondary (Alexa 488, 1:1000) antibodies and mounted on charged slides (Superfrost plus; Fisher Scientific) with Vectasheild containing DAPI. Images were obtained with a Leica DM6 epifluorescence microscope. Brightness and contrast were adjusted manually using Adobe Photoshop; z-stacks were generated using Leica software.

#### Volumetric Analysis

Quantification of cortical lesion volume was performed by measuring the area of cortical tissue remaining in both hemispheres in eight DAPI-labeled coronal sections along ~2400 μm of the rostral-caudal axis spaced 300 μm apart as previously described (Frankowski et al., 2019; Zhu et al., 2019). Borders of the cortical plate were drawn between the dorsal aspect of the corpus callosum and the pial surface using ImageJ. Regions of the cortical subplate (e.g., amygdala) were excluded from analysis. The % of the ipsilateral cortex remaining for each animal was calculated using the following formula:

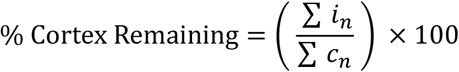

where i = the area of the ipsilateral cortex and c = the area of the contralateral cortex and n = the section number.

#### Cortical thickness measurement

Average cortical thickness was measured from a series of three DAPI-labeled x10 images of the entire cortical column centered at the injury epicenter and two 300 μm serial sections caudal to the epicenter. The area of tissue between the pial surface and the ventral aspect of layer 5/6 divided by the width of the frame (958.29 μm) to obtain an average cortical thickness value along the width of the frame. For uninjured-controls, images were taken in corresponding brain sections at the most central portion of V1 as defined in the 2017 Allen Reference Atlas.

#### Cell quantification

Fluorescently labeled coronal brain sections were imaged using a Leica DM6 fluorescence microscope with an x10 objective and quantification was performed in ImageJ, as previously described (Frankowski et al., 2019; Zhu et al., 2019). For quantification of GFP+ cell density, three brain sections spaced 300 μm apart were counted, with the rostral-most section at the injury epicenter and the next two additional sections caudal to the epicenter.

#### Neurophysiology

Animals were initially anesthetized with 2% isoflurane in a mixture of N2_O_/O_2_ (70%/30%) then placed into a stereotaxic apparatus. A small, custom-made plastic chamber was glued (Vetbond, St. Paul, MN, US) to the exposed skull. After one day of recovery, re-anesthetized animals were placed in a custom-made hammock, maintained under isoflurane anesthesia (1-2% in N_2_O/O_2_) and multiple single tungsten electrodes were inserted into V1 using the same craniotomy performed during the injury phase. Following electrodes placement, the chamber was filled with sterile agar and sealed with sterile bone wax. Animals were then sedated with chlorprothixene hydrochloride (1 mg/kg; IM; (Camillo et al., 2018)) and kept under light isoflurane anesthesia (0.2 – 0.4% in 30% O_2_) throughout the recording procedure. EEG and EKG were monitored throughout and body temperature was maintained with a heating pad (Harvard Apparatus, Holliston, MA).

Data was acquired using a multi-channel Scout recording system (Ripple, UT, USA). Local field potentials (LFP) from multiple locations were band-pass filtered from 0.1 Hz to 250 Hz and stored along with spiking data at 1 kHz sampling rate. LFP signal was aligned to stimulus time stamps and averaged across trials for each recording location in order to calculate visually evoked potentials (VEP)(Foik et al., 2015; Suh et al., 2020). Single neuron spike signals were band-pass filtered from 500 Hz to 7 kHz and stored at a 30 kHz sampling frequency. Spikes were sorted online in Trellis (Ripple, UT, USA) while performing visual stimulation. Visual stimuli were generated in Matlab (Mathworks, USA) using Psychophysics Toolbox (Brainard, 1997; Pelli, 1997; Kleiner et al., 2007) and displayed on a gamma-corrected LCD monitor (55 inches, 60 Hz; 1920 × 1080 pixels; 52 cd/m^2^ mean luminance). Stimulus onset times were corrected for monitor delay using an in-house designed photodiode system (Foik et al., 2018).

Visual responses were assessed using methods similar to previous publications (Foik et al., 2018; Foik et al., 2020; Suh et al., 2020). For recordings of visually evoked responses, cells were first tested with 100 repetitions of a 500 ms bright flash stimulus (105 cd/m^2^). Receptive fields for visually responsive cells were then located using square-wave drifting gratings, after which optimal orientation, direction, and spatial and temporal frequencies were determined using sine wave gratings. Spatial frequencies used ranged from 0.001 to 0.5 cycles/º; Temporal frequencies used were from 0.1 to 10 cycles/s. Using optimal parameters, size tuning was assessed with apertures ranging from 1 to 110º at 100% contrast. With optimal size, orientation tuning of the cell was re-assessed using 8 orientations × 2 directions each, stepped by 22.5º increments.

#### Local Field Potential (LFP) Analysis

LFP signal was normalized using z-score standardization as we have done previously (Foik et al., 2015; Wypych et al., 2012, 2014). Amplitude of response was calculated as a difference between the peak of the positive and negative components of the VEP. Response latency was defined as the time point of maximum response. Maximum of the response was defined at the larger of the negative or positive peak.

#### Single Unit Analysis

Tuning curves were calculated based on average spike rate. Optimal visual parameters were chosen as the maximum response value. Orientation tuning was measured in degrees at the half-width at half-height (HWHH; 1.18 × σ) based on fits to Gaussian distributions (Alitto and Usrey, 2004; Carandini and Ferster, 2000; Foik et al., 2018, 2020; Liu et al., 2011, 2015) using:

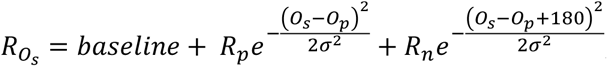

where O_s_ is the stimulus orientation, R_Os_ is the response to different orientations, O_p_ is the preferred orientation, R_p_ and R_n_ are the responses at the preferred and non-preferred direction, σ is the tuning width, and ‘baseline’ is the offset of the Gaussian distribution. Gaussian fits were estimated without subtracting spontaneous activity, similar to the procedures of Alitto and Usrey (Alitto and Usrey, 2004).

Size tuning curves were fitted by a difference of Gaussian (DoG) function:

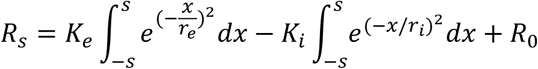

in which R_s_ is the response evoked by different aperture sizes. The free parameters, K_e_ and re, describe the strength and the size of the excitatory space, respectively; Ki and ri represent the strength and the size of the inhibitory space, respectively; and R_0_ is the spontaneous activity of the cell.

The optimal spatial and temporal frequency was extracted from the data fitted to Gaussian distributions using the following equation (DeAngelis et al., 1993; Van Den Bergh et al., 2010; Foik et al., 2018, 2020):

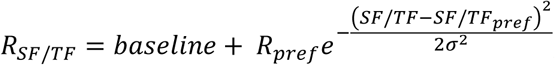

Where R_SF/TF_ is the estimated response, R_pref_ indicates response at preferred spatial or temporal frequency, SF/TF indicates spatial or temporal frequency, σ is the standard deviation of the Gaussian, and baseline is Gaussian offset.

### Quantification and Statistical Analysis

Anatomical data analysis was performed in Graphpad Prism 6 and Microsoft Excel. Experimental groups were compared by unpaired two-tailed t-test, two-way ANOVA with Tukey’s post-hoc tests, or repeated measures two-way ANOVA followed by Sidak’s post-hoc test. Neurophysiology data analysis was performed in Matlab (Mathworks, USA). Average differences between groups were compared using two-tailed Mann-Whitney U tests. All data are expressed as mean ± SEM. Significance was set at P < 0.05.

### Data and Software Availability

All data that support the findings of this study are available from the corresponding authors upon reasonable request.

---

**Figure S1.**
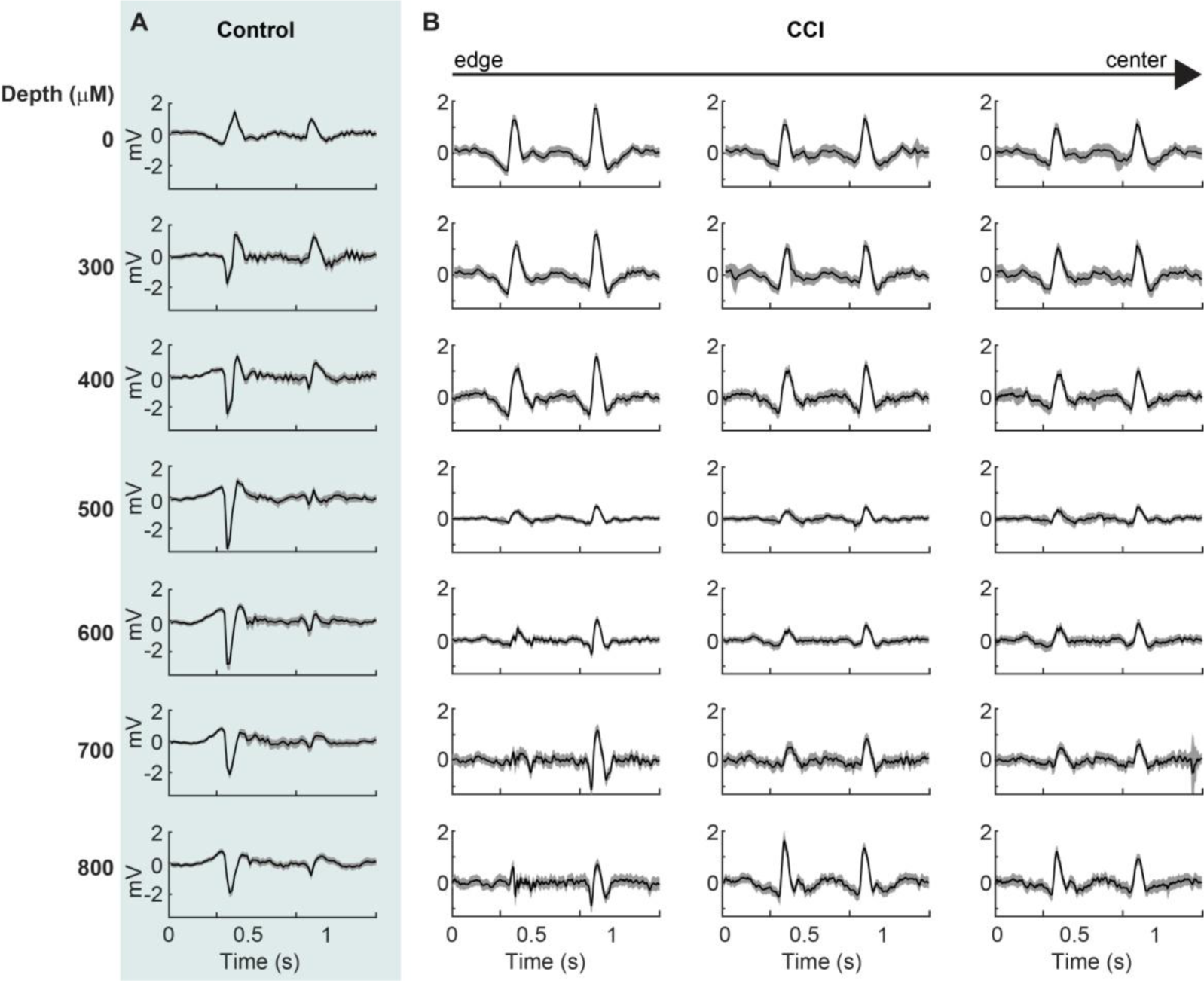
Visual response profile in V1 is severely degraded by TBI. **A.** In an uninjured control animal, the cortical VEP profile starts from large positive components indicating output from layer II/III that (0 – 400 μm) reverses into negative components in layers IV, V, and VI (500 – 1,000 μm). **B.** After CCI, from the edge to the center of the injury site (arrow), the cortical profile is abnormal compared to control. Amplitudes are smaller, particularly at 500 μm and deeper, and there is an absence of a robust negative wave component. Black lines represent average response to flash of light and grey shading indicates SEM.

